# Signaling bias of the protease-activated receptor-1 is dictated by distinct GRK5 and β-arrestin-2 determinants

**DOI:** 10.1101/2025.11.02.686167

**Authors:** Monica L. Gonzalez Ramirez, Lennis B. Orduña-Castillo, Carolyne Bardeleben, Huaping Qin, Ying Lin, Cierra A. Birch, Irina Kufareva, JoAnn Trejo

**Affiliations:** Department of Pharmacology, School of Medicine, University of California, San Diego, La Jolla, CA 92093, USA; Skaggs School of Pharmacy and Pharmaceutical Sciences, University of California, San Diego, La Jolla, CA 92093, USA

**Keywords:** APC, activated protein C, biased agonists, biased signaling, G protein, GPCR, G protein-coupled receptor, kinase, PAR1, phosphorylation, thrombin

## Abstract

G protein-coupled receptors (GPCRs) exhibit signaling bias, or preferential activation of heterotrimeric G proteins versus GPCR receptor kinase (GRK)-mediated β-arrestin signaling. The protease-activated receptor-1 (PAR1) GPCR activates both G protein and β-arrestin in response to thrombin, but only β-arrestin in response to activated protein C (APC). Thrombin-activated PAR1-G protein signaling is desensitized by β-arrestin-1, whereas APC-activated PAR1 signaling is propagated by β-arrestin-2. The mechanisms underlying PAR1 biased signaling are not known. Here, using computational modeling and cell biology studies, we reveal the molecular basis of signaling by thrombin- and APC-activated PAR1. Although both thrombin- and APC-induced PAR1 signaling are regulated by the same GRK isoform, GRK5, the two types of signaling are differentially dependent on GRK5 membrane anchoring, PAR1 C-terminal phosphorylation, and the binding mode of β-arrestin-2. These differences translate into distinct β-arrestin-2 conformations and define the cytoprotective signaling signature by APC which contrasts with thrombin inflammatory signaling.

## Introduction

G protein-coupled receptors (GPCRs) transduce signals in response to a wide range of stimuli, control vast physiological functions and are highly druggable. Upon agonist binding, GPCRs undergo conformational changes that facilitate activation of heterotrimeric G protein signaling. Agonist-activated GPCRs are phosphorylated by GPCR kinases (GRKs), which enables β-arrestin recruitment. β-arrestins desensitize agonist-activated GPCR-G protein signaling and function as scaffolds that assemble and activate signaling complexes. GPCRs display bias toward either G protein or GRK-mediated β-arrestin signaling depending on the activating agonist ^1, 2^. However, the mechanisms that specify how biased agonists induce GPCR preferential activation of heterotrimeric G proteins *versus* β-arrestin-mediated signaling are complex and not well understood.

Protease-activated receptor-1 (PAR1) is a GPCR for the coagulant protease thrombin ^3^ and the anti-coagulant protease activated protein C (APC) ^4^ and an important drug target ^5^. Thrombin-activation of PAR1 promotes G protein signaling and mediates platelet activation leading to thrombosis ^6, 7^, cancer progression ^8^ and endothelial inflammatory responses including p38 mitogen activated protein kinase (MAPK) signaling ^9, 10^. The PAR1 antagonist vorapaxar is approved for thrombotic cardiovascular events ^7^. In contrast to thrombin, APC-activated PAR1 promotes β-arrestin-2 (βarr2)-mediated cytoprotective responses including endothelial barrier stabilization, anti-apoptotic Akt pro-survival activities ^11, 12^ and βarr2-dependent neuroprotection in a mouse model ^13^, making APC variants or synthetic ligands promising therapeutics. Thrombin and APC biased agonism is mediated by differential cleavage of the PAR1 N-terminus resulting in the generation of distinct tethered activating ligands. Thrombin cleaves PAR1 at an N-terminal arginine (R)-41 site, generating a tethered ligand that triggers G protein signaling ^6^ and is rapidly desensitized by β-arrestin-1 (βarr1) ^14, 15^. By contrast APC bound to the endothelial protein C receptor (EPCR), a transmembrane cofactor, cleaves PAR1 at an N-terminal R-46 site generating a different tethered ligand and activates βarr2 cytoprotective signaling ^11, 12, 16, 17^.

Structural studies indicate that adoption of distinct conformational states by GPCRs, GRKs and β-arrestins is critical for specifying biased signaling ^1, 18, 19^. The pattern or barcode of phosphorylation sites on GPCRs induced by different GRKs in response to biased agonists are known to differently modulate β-arrestin function ^18, 20, 21^. The GRKs 2,3,5 and 6 isoforms are widely expressed, however, the mechanisms that specify which individual GRK isoform regulates specific GPCR-stimulated β-arrestin functions in response to biased agonists is not well understood. Here we sought to understand the molecular basis by which GRKs and β-arrestins drive PAR1 biased signaling.

In this study we report that thrombin- and APC-mediated activation of PAR1 promotes distinct conformational states of the receptor transmembrane (TM) domain. However, thrombin- and APC-induced PAR1 signaling are both regulated by the same GRK isoform, GRK5. Despite this, the two types of PAR1 biased signaling rely on different GRK5 and βarr2 determinants, distinct receptor C-terminus phosphorylation patterns, and different modes of βarr2 engagement. This translates into distinct conformations of βarr2 that may mediate the proinflammatory *versus* cytoprotective signaling induced by thrombin and APC activation of PAR1.

## Results

### Structural predictions of PAR1 conformational states induced by biased agonists

The crystal structure of PAR1 bound to the antagonist vorapaxar ^22^ and recent structures of activated PAR1 bound to the thrombin-generated tethered activating ligand have been reported ^23, 24^. However, the structure of APC-activated PAR1 is not known. To gain insight into the conformational preferences of unactivated PAR1, thrombin-activated PAR1, and APC-activated PAR1, we built an ensemble of 100 structural models of each of these molecular species using AlphaFold 3 (AF3) ^25^. To reduce conformational bias, all models were built without intracellular effectors or the disordered C-terminus and included amino acid residues 22-390, 42-390, and 47-390 for unactivated PAR1, thrombin-activated PAR1, and APC-activated PAR1, respectively. The model ensembles highlighted clear conformational differences (Fig. 1a-c). Thrombin-activated PAR1 showed high conformational consistency across the ensemble, with the N-terminal tethered ligand docking into the shallow and narrow orthosteric binding site and interacting with key residues important for activation (Fig. 1b), as recently reported ^23, 24^. In contrast, the ensembles of the unactivated PAR1 and APC-activated PAR1 displayed substantial conformational variability (Fig. 1a, c). The full-length N-terminus of uncleaved PAR1 was predicted to partially dock in the orthosteric pocket in some models and stayed out in the solvent in others (Fig. 1a). The APC-cleaved N-terminus invariably stayed out of the pocket in all 100 models (Fig. 1c).

**Fig. 1.**
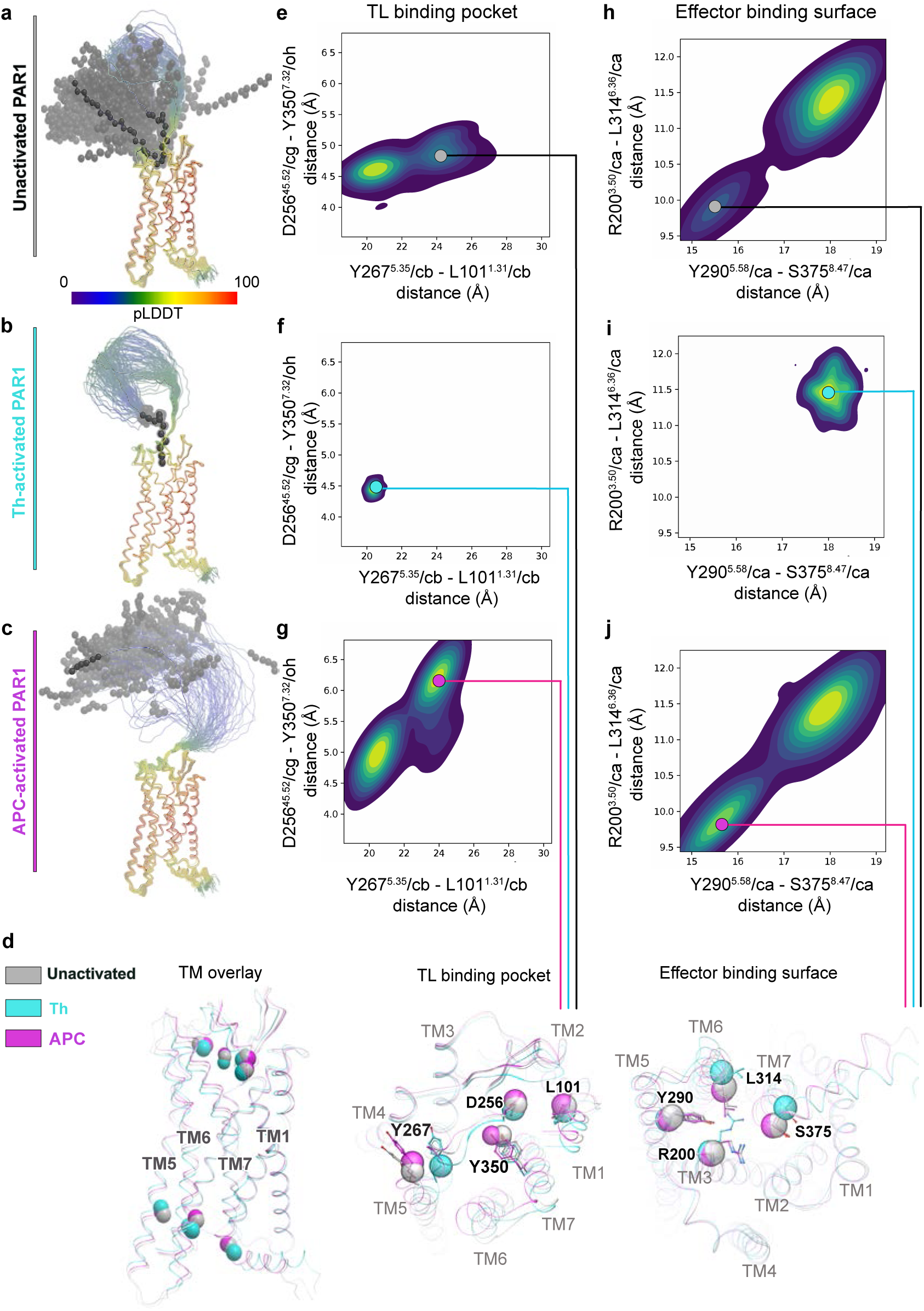
Predicted structures and conformational preferences of unactivated, thrombin-activated and APC-activated PAR1. Structural ensembles of 100 AlphaFold 3 models of unactivated PAR1 (**a**), thrombin-activated PAR1 (**b**) and APC-activated PAR1 (**c**). Receptors are shown as backbone ribbons and colored by the per-residue pLDDT score (reflecting prediction confidence). The distributions, across the model ensemble, of distances between key residues in the orthosteric tethered ligand (TL) binding pocket (*middle*) and the effector binding surface (*right*) are shown as pseudo-colored two-dimensional density plots. Distances are measured in angstroms (Å). Plots, left-to-right, represent the ensembles of unactivated PAR1 (**e, h**), thrombin-activated PAR1 (**f, i**), and APC-activated PAR1 (**g, j**). Small, filled circles mark representative conformations specific for each N-terminally cleaved and uncleaved species and shown in (**d**). **d**, overlay of representative models for unactivated (*silver*), thrombin-activated (*cyan*) and APC-activated (*magenta*) PAR1 transmembranes (TMs). Receptor molecules are shown in ribbon with atoms used for calculating the distances shown as sticks and spheres in the transmembrane overlay (*left*). The *middle* panel is a view across the membrane plane from the extracellular side, showing the TL binding pocket, while the *right* panel is a view from the intracellular side, focusing on the effector binding surface.

To understand differences in the preferred conformations of PAR1 transmembrane (TM) helical bundle, corresponding to differentially cleaved receptor species, we measured distances between key residues in the PAR1 tethered ligand binding pocket (D256^45.52^/cg – Y350^7.32^/oh and Y267^5.35^/cb-L101^1.31^/cb) and the effector binding surface (R200^3.50^/ca – L314^6.36^/ca and Y290^5.58^/ca – S375^8.47^/ca). In contrast to unactivated PAR1 (Fig. 1d,e), the thrombin-activated PAR1 distance map exhibited a constricted tethered ligand (TL) binding pocket with shorter distances between the key residue pairs D256-Y350 and Y267-L101 (Fig. 1d,f) and a uniform extended effector binding surface with longer distances between key residues R200-L314 and Y290-S375 (Fig. 1d, i). Similar to other Class A GPCRs ^26^, this signature corresponds to an active state of PAR1 that favors coupling to heterotrimeric G proteins. By contrast, the orthosteric binding pocket distance maps in the APC-activated PAR1 demonstrated more variable and frequently more open ligand binding pocket conformations (Fig. 1d,g), compared to thrombin-activated PAR1 (Fig. 1d,f) and unactivated PAR1 (Fig. 1d,e). Consistent with the idea of allosteric communication between the extracellular orthosteric and the intracellular effector-binding pockets, APC-activated PAR1 preferentially featured a narrow effector binding surface (Fig. 1d,j), which was not observed in thrombin-activated PAR and only rarely observed in unactivated PAR1 (Fig. 1h,i). The narrow effector binding interface in APC-activated PAR1 makes it incompatible with the canonical G protein coupling geometry and explains the experimentally observed bias of APC-activated PAR1 towards βarr2 ^11, 12, 13^. Together, these findings suggest that thrombin- and APC-activated PAR1 display distinct active conformational states that likely mediate the balanced PAR1-G-protein-βarr1 coupling induced by thrombin and βarr2-biased signaling promoted by APC.

### A dual role for GRK5 in regulation of protease-activated receptor-1 signaling bias

To delineate the function of GRKs in PAR1 biased signaling, we first examined GRK2, GRK3, GRK5 and GRK6 mRNA transcript abundance in human cultured endothelial cells using RT-qPCR. In human umbilical vein endothelial cells (HUVEC)-derived EA.hy926 cells, GRK5 showed the highest abundance, whereas GRK2 and GRK6 mRNA transcript expression were detected at ∼50% and ∼25% of GRK5 mRNA, respectively (Fig. 2a). Expression of GRK3 mRNA transcripts was nearly negligible in endothelial EA.hy926 cells (Fig. 2a). A similar profile of GRK2,3,5 and 6 expression was detected in primary HUVECs (Fig. 2b), indicating that GRK5 and GRK2 are the most abundant GRKs expressed in human cultured endothelial cells and are more likely to play a role in PAR1 biased signaling.

**Fig. 2.**
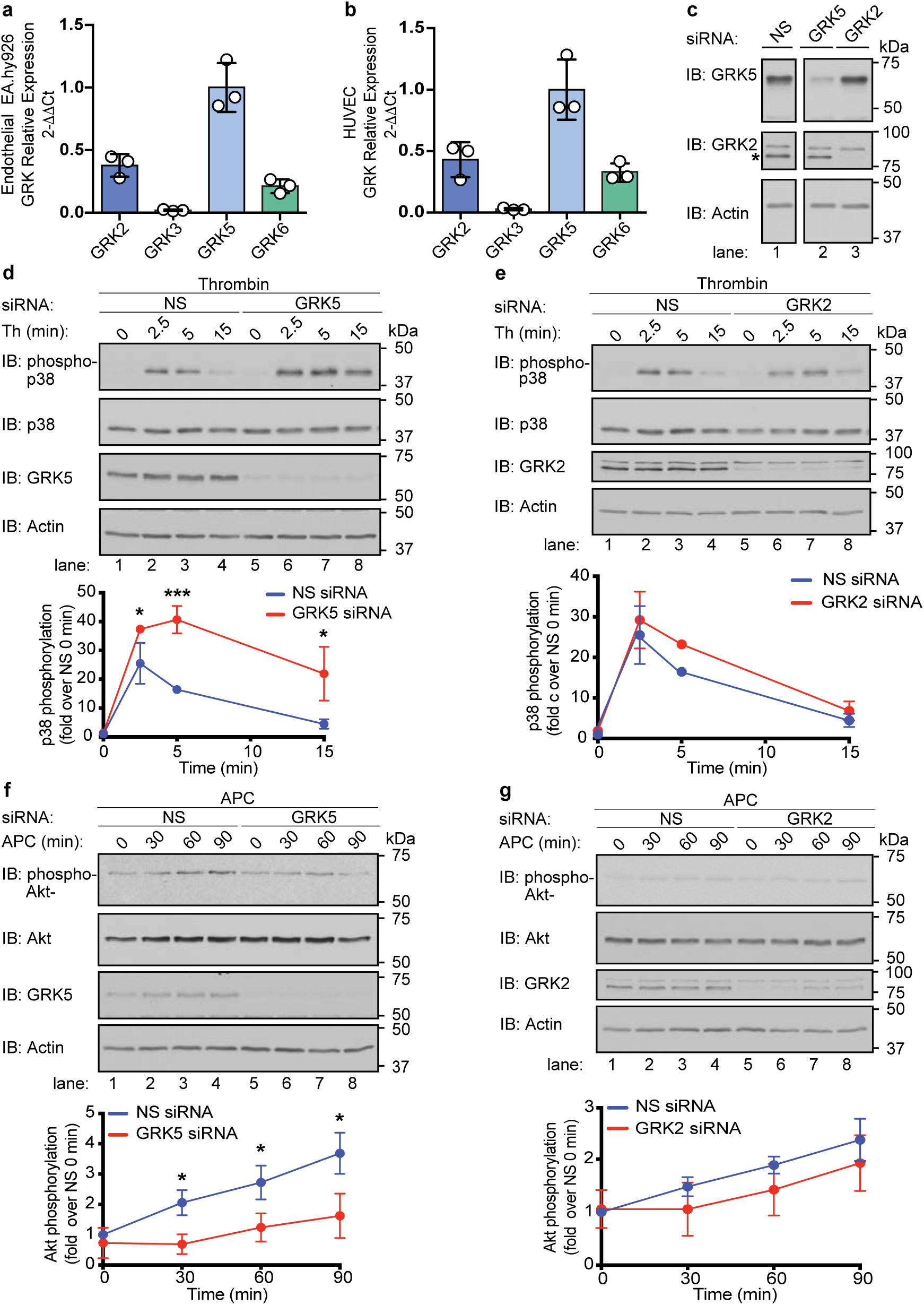
GRK5 regulates both thrombin and APC biased signaling. Relative GRK2, GRK3, GRK5, and GRK6 mRNA transcript expression in (**a**) HUVEC-derived endothelial EAhy.926 cells and (**b**) primary HUVEC were quantified by RT-qPCR and normalized to 18S ribosomal mRNA expression. The data (mean ± S.D.) from three independent experiments is expressed as a fraction of GRK5 mRNA transcript expression. **c**, Cell lysates from EA.hy926 cells transfected with GRK2, GRK5 or non-specific (NS) siRNA were immunoblotted to detect endogenous GRK5 and GRK2 (asterisk). Actin expression was used as a loading control. Endothelial EA.hy926 cells transfected with NS or GRK5 siRNA (**d** and **f**) or NS and GRK2 siRNA (**e** and **g**) were incubated with either 10 nM thrombin for 0 min to 15 min or 20 nM APC for 0 min to 90 min. Cell lysates were immunoblotted as indicated. The data (mean ± S.D.) from three independent experiments are expressed as fold change over NS siRNA at 0 min. Statistical significance was determined by two-way ANOVA followed by Šídák’s multiple comparisons test (*, *p* < 0.05; ***, *p* < 0.001).

To directly assess GRK5 and GRK2 function in PAR1 biased signaling, an siRNA knockdown approach was used in human cultured endothelial cells. In control cells transfected with control non-specific siRNA, GRK5 was detected as an ∼65 kDa protein and GRK2 migrated as an ∼80 kDa species (Fig. 2c, lane 1). The GRK5-specific siRNA markedly reduced expression of endogenous GRK5 but not GRK2 (Fig. 2c, lanes 2 vs. 3), whereas the GRK2-specific siRNA virtually abolished GRK2 expression without altering GRK5 expression (Fig. 2c, lanes 3 vs. 2). To assess the effect of GRK5 and GRK2 knockdown on thrombin signaling, we examined p38 MAPK activation, which is mediated by G protein signaling downstream of activated PAR1^27^. In endothelial cells transfected with non-specific control siRNA, thrombin induced a marked increase in p38 MAPK T180/Y182 phosphorylation at 2.5 min that declined by 15 min (Fig. 2d, lanes 1-4). In GRK5 depleted endothelial cells, p38 phosphorylation was significantly increased and prolonged after thrombin treatment compared to control cells (Fig. 2d). In contrast, the knockdown of GRK2 expression failed to alter thrombin-stimulated p38 phosphorylation (Fig. 2e, lanes 1-8).

To determine the function of GRK5 and GRK2 in the biased signaling of APC-activated PAR1, we examined Akt pro-survival signaling which is mediated by βarr2 in endothelial cells ^12,28^. In non-specific control siRNA transfected cells, APC induced a significant increase in Akt S473 phosphorylation at 30 min that remained elevated for 90 min (Fig. 2f, lanes 1-4), consistent with previous reports ^12^. However, in GRK5-depleted cells, APC-stimulated Akt signaling was virtually abolished (Fig. 2f, lanes 5-8), whereas siRNA knockdown of GRK2 expression had no effect on APC-stimulated Akt signaling (Fig. 2g, lanes 1-8). These data indicate a critical role for GRK5 in regulating both thrombin- and APC-activated PAR1 signaling.

### GRK5 is required for both thrombin- and APC-induced βarr2 recruitment

To further interrogate GRK5 function in PAR1 signaling and bias, we optimized a bioluminescence resonance energy transfer (BRET) system in HEK293 cells to recapitulate thrombin and APC-induced β-arrestin recruitment. APC requires the EPCR cofactor to facilitate cleavage and activation of PAR1 ^29, 30^. Therefore, we used HEK293 cells expressing PAR1-fused to yellow fluorescent protein (YFP) and Renilla luciferase (Rluc)-βarr2 either with or without EPCR co-expression. Cells were stimulated with thrombin or APC, respectively and βarr2 recruitment was assessed by BRET (Fig. 3a). Thrombin induced a rapid and significant increase in βarr2 recruitment to PAR1 regardless of EPCR expression (Fig. 3b) and this increase was completely blocked by vorapaxar (Fig. 3b). APC induced a hyperbolic and significant increase in PAR1-βarr2 net BRET in cells co-expressing EPCR, which was also abolished by vorapaxar (Fig. 3c). In cells lacking EPCR expression, APC-stimulated PAR1-induced βarr2 recruitment yielded a quasi-linear net BRET increase, suggestive of random collisions rather than specific interactions (Fig. 3c) and supporting the notion that EPCR is required for APC efficient activation of PAR1.

**Fig. 3.**
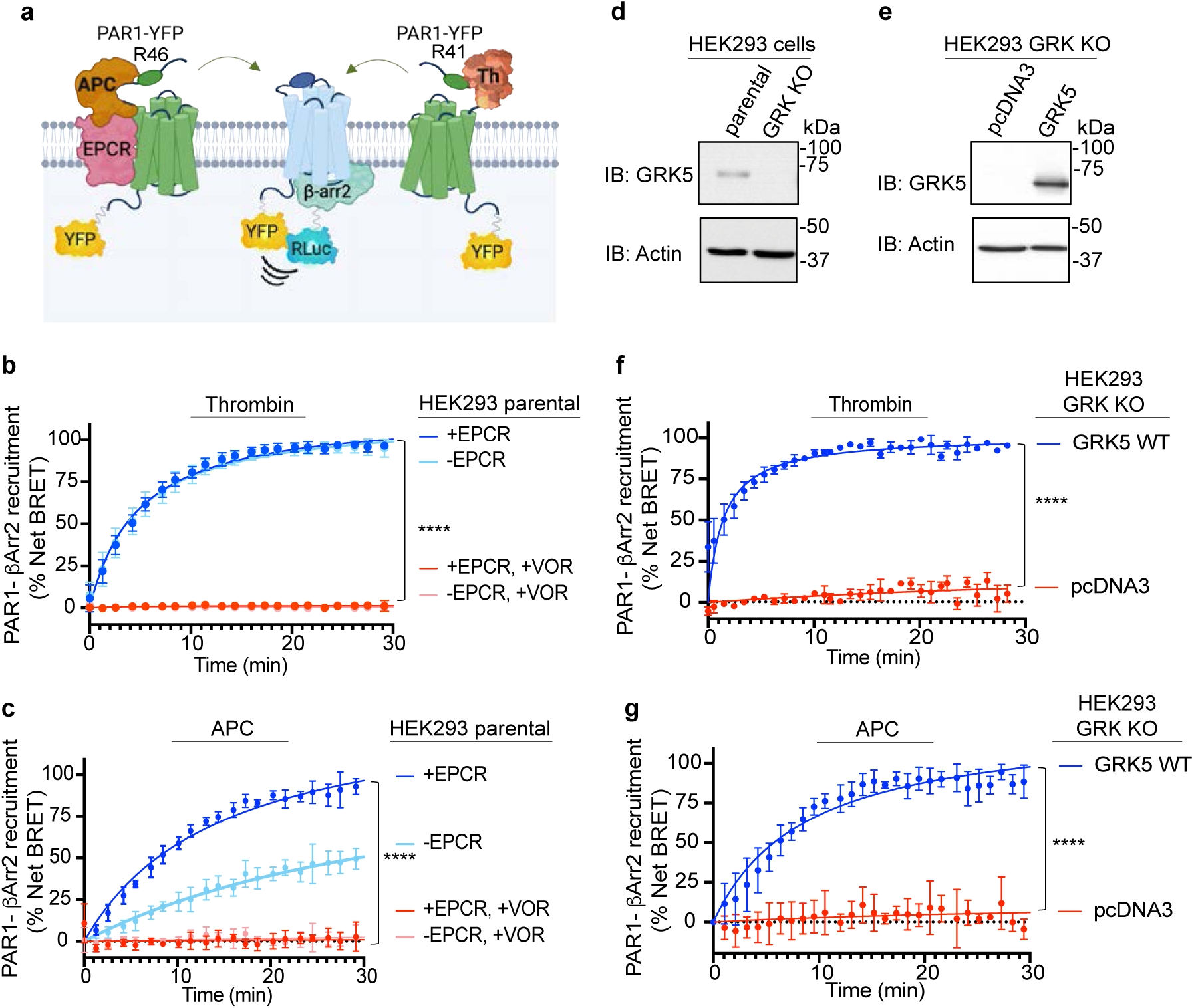
GRK5 is both necessary and sufficient for APC- and Th-activated PAR1 induced βarr2 recruitment. **a**, Cartoon of BRET assay showing recruitment of Rluc-βarr2 induced by either APC-EPCR activation of PAR1-YFP through arginine (R)46 cleavage or thrombin-activation of PAR1-YFP via R41 cleavage. HEK293 cells transfected with PAR1-YFP, EPCR-Halo and Rluc-βarr2, pretreated with vorapaxar or DMSO vehicle were stimulated with 1 nM thrombin (**b**) or 20 nM APC (**c**) and βarr2 recruitment to PAR1 was determined by BRET. The data (mean ± S.D.) from three independent repetitions was analyzed by one-way ANOVA followed by Tukey’s multiple comparisons test (****, *p* < 0.0001). **d** and **e**, cell lysates from HEK293 parental cells and HEK293 CRISPR-Cas9 GRK knockout (KO) cells transfected with GRK5 wildtype (WT) or pcDNA3 vector were immunoblotted as indicated. HEK293 GRK KO cells transfected with PAR1-YFP, EPCR-Halo and Rluc-βarr2 co-expressing GRK5 WT or pcDNA3 were stimulated with 1 nM thrombin (**f**) or 20 nM APC (**g**) and recruitment of βarr2 to PAR1 was determined by BRET. The data (mean ± S.D.) from three independent replicates was analyzed by *t* test (****, *p* < 0.0001).

We next examined whether GRK5 expression was required for thrombin- and APC-activated PAR1 recruitment of βarr2 using HEK293 quadruple GRK2,3,5,6 CRISPR-Cas9 knockout (KO) cells ^31^ with and without re-expression of wildtype GRK5 (Fig. 3d, e). This system provides an effective strategy to dissect the role of GRK5 in PAR1 signaling and bias, avoids potential contributions from other GRKs, and minimizes clone-associated issues ^31, 32^. In HEK293 GRK KO cells expressing PAR1-YFP, Rluc-βarr2, and EPCR, no specific increase in PAR1-βarr2 BRET was observed upon thrombin stimulation; however, the signal was fully rescued by re-expression of GRK5 (Fig. 3e, f). Similarly, re-expression of GRK5 restored APC-activated PAR1-induced βarr2 recruitment (Fig. 3e, g). Thus, GRK5 expression is both necessary and sufficient for recruitment of βarr2 to PAR1 following activation with thrombin or APC.

### Distinct GRK5 determinants specify thrombin-*versus* APC-induced βarr2 recruitment

GRK5 localizes to the plasma membrane via a C-terminal amphipathic helix, associates with agonist-activated GPCRs, and promotes receptor phosphorylation ^33^. To determine whether specific GRK5 determinants regulate βarr2 recruitment to thrombin-and APC-activated PAR1, we utilized two GRK5 mutants. The first mutant, termed 4A ^34^, features alanine (A) substitutions for leucine (L550, L551, L554) and phenylalanine (F555) residues of the amphipathic helix, and is defective in plasma membrane localization. The second mutant features a lysine (K)215 to arginine (R) substitution in the active site and is catalytically inactive. The localization of GRK5 WT, 4A and K215R mutants in transiently transfected HeLa cells was confirmed by immunofluorescence confocal microscopy. Confocal imaging combined with line-scan analysis indicated that GRK5 WT and the catalytically inactive K215R mutant reside primarily at the plasma membrane (Fig. 4a), whereas the GRK5 4A mutant redistributed predominantly to the cytoplasm (Fig. 4a), as previously reported ^34^.

**Fig. 4.**
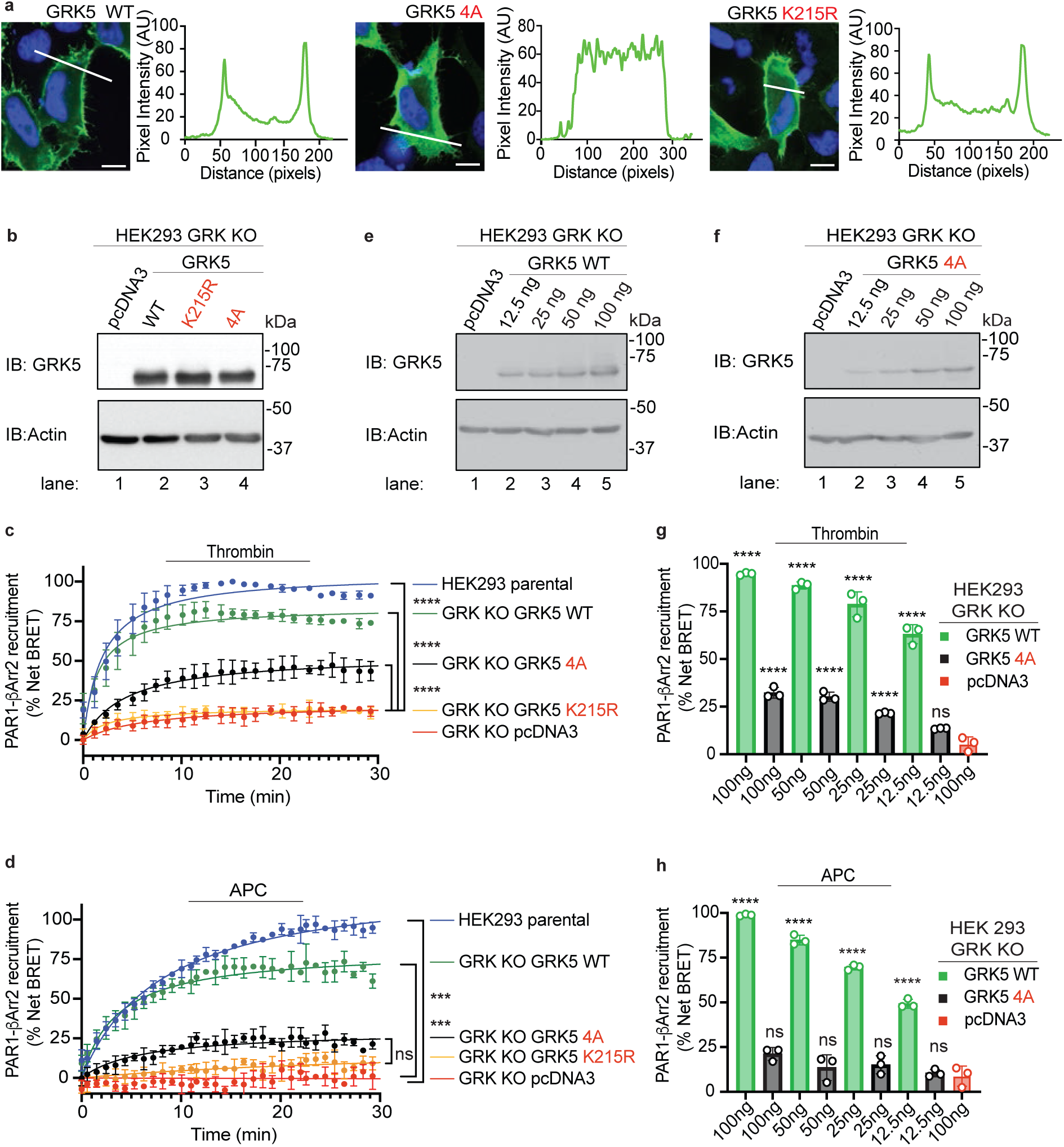
Distinct GRK5 determinants regulate APC-*versus* Th-induced βarr2 recruitment to PAR1. **a**, Subcellular localization of GRK5 wildtype, K215R and 4A mutant (*green*) was verified by immunofluorescence confocal microscopy using HeLa cells. Nucleus is stained with DAPI. Line scan analysis was performed in Image J. Scale bar, 100 μm. **b**, HEK293 CRISPR-Cas9 GRK knockout (KO) cells transfected with GRK5 wildtype (WT), K215R, 4A mutant or pcDNA3 were immunoblotted as indicated. Actin was used as a loading control. **c** and **d**, HEK293 GRK KO cells co-expressing PAR1-YFP, EPCR-Halo and Rluc-βarr2 with either GRK5 WT, K215R, 4A mutant or pcDNA3 were stimulated with 1 nM thrombin or 20 nM APC and βarr2 recruitment to PAR1 was determined by BRET. HEK293 parental WT cells were treated similarly and recruitment of βarr2 determined by BRET. The data (mean ± S.D.) from three independent replicates was analyzed by one-way ANOVA followed by Tukey’s multiple comparisons test (***, *p* < 0.001; ****, *p* < 0.0001; ns, not significant). **e** and **f,** HEK293 GRK KO cell lysates transfected with increasing amounts of GRK5 WT and 4A mutant or pcDNA3 were immunoblotted as shown. **g** and **h**, HEK293 GRK KO cells co-expressing PAR1-YFP, EPCR-Halo, Rluc-βarr2 and increasing amounts of GRK5 WT and 4A mutant or pcDNA3 were stimulated with 1 nM thrombin or 20 nM APC and recruitment of βarr2 to PAR1 determined by BRET. The data (mean ± S.D.) from three independent repetitions was analyzed by one-way ANOVA followed by Tukey’s multiple comparisons test (****, *p* < 0.0001; ns, not significant).

The effect of GRK5 WT and mutants on thrombin- and APC-activated PAR1-induced βarr2 recruitment was next examined using BRET and HEK293 GRK KO cells. HEK293 GRK KO cells expressing PAR1-YFP, Rluc-βarr2, and EPCR were co-transfected with equivalent amounts of either GRK5 WT, K215R, 4A mutant or pcDNA3 (Fig. 4b). In both thrombin- and APC-stimulated cells, βarr2 recruitment to PAR1 was abolished in GRK KO cells expressing the K215R catalytic mutant or pcDNA3 and restored in cells re-expressing GRK5 WT comparable to that observed in HEK293 parental cells (Fig. 4c, d). However, in cells re-expressing the GRK5 4A mutant at levels comparable to GRK5 WT expression (Fig. 4b), APC/PAR1-induced βarr2 recruitment was negligible and not significant compared to the catalytic inactive K215R mutant (Fig. 4d), indicating that both GRK5 membrane localization and catalytic activity are critical for βarr2 recruitment to APC-activated PAR1. By contrast, re-expression of GRK5 4A mutant resulted in a significant increase in βarr2 recruitment to thrombin-activated PAR1 (Fig. 4c), although such recruitment was still lower than that mediated by GRK5 WT. As expected, the catalytically inactive GRK5 K215R mutant failed to restore thrombin-induced βarr2 recruitment (Fig. 4c), indicating that the catalytic activity of GRK5 is essential for βarr2 recruitment to PAR1 in response to both thrombin and APC.

To further examine the impact of GRK5 WT and 4A mutant on PAR1 signaling and bias, we examined the effect of different GRK5 WT and 4A mutant expression levels on βarr2 recruitment to thrombin- and APC-activated PAR1 by BRET. Compared to βarr2 recruitment to thrombin-activated PAR1 in HEK293 GRK KO cells re-expressing GRK5 WT, the lowest level of GRK5 WT expression allowed for ∼60% of βarr2 recruitment signal (Fig. 4e, g). Similarly, increasing expression of GRK5 WT resulted in a concomitant increase in APC-induced βarr2 recruitment with the lowest level of GRK5 WT expression resulting in an ∼50% increase in βarr2 recruitment to APC-activated PAR1 (Fig. 4e, h). GRK4A mutant expression resulted in a partial but significant increase in thrombin-stimulated βarr2 recruitment at all except the lowest expression levels (Fig. 4f, g). In contrast to GRK5 WT, GRK4A mutant failed to significantly enhance APC-induced βarr2 recruitment even at the highest level of expression (Fig. 4f, h). Thus, unlike thrombin, APC-induced βarr2 recruitment is critically dependent on GRK5 membrane localization for ýarr2 recruitment.

### Thrombin and APC require distinct PAR1 C-terminal phosphorylation sites for GRK5-dependent βarr2 recruitment

GPCR biased signaling is driven in part by distinct patterns of GRK-mediated phosphorylation of the C-terminus. The C-terminus sequence of PAR1 contains five serine phosphorylated residues pS396, pS399, pS400, pS412 and pS418 that have been previously identified to be phosphorylated by mass spectrometry (Fig. 5a, b) ^35^. A number of additional Ser and Thr sites of phosphorylation in the C-terminus have also been reported ^35^. To determine if different PAR1 phosphorylation sites are required for thrombin- and APC-induced βarr2 recruitment, three PAR1 C-terminal mutants were generated. First was a fully phospho-deficient PAR1 0P mutant where all serine (S) and threonine (T) residues within the receptor’s helix 8 and the C-terminus were mutated to alanine (A). In the dP2 mutant, all candidate Ser/Thr phosphorylation sites in the proximal C-terminus were mutated to alanine while those in the distal C-terminus were retained. In the dP3 mutant, candidate Ser/Thr phosphorylation sites in the distal C-terminus were mutated to alanine while those in the proximal C-terminus were retained (Fig. 5a, b).

**Fig. 5.**
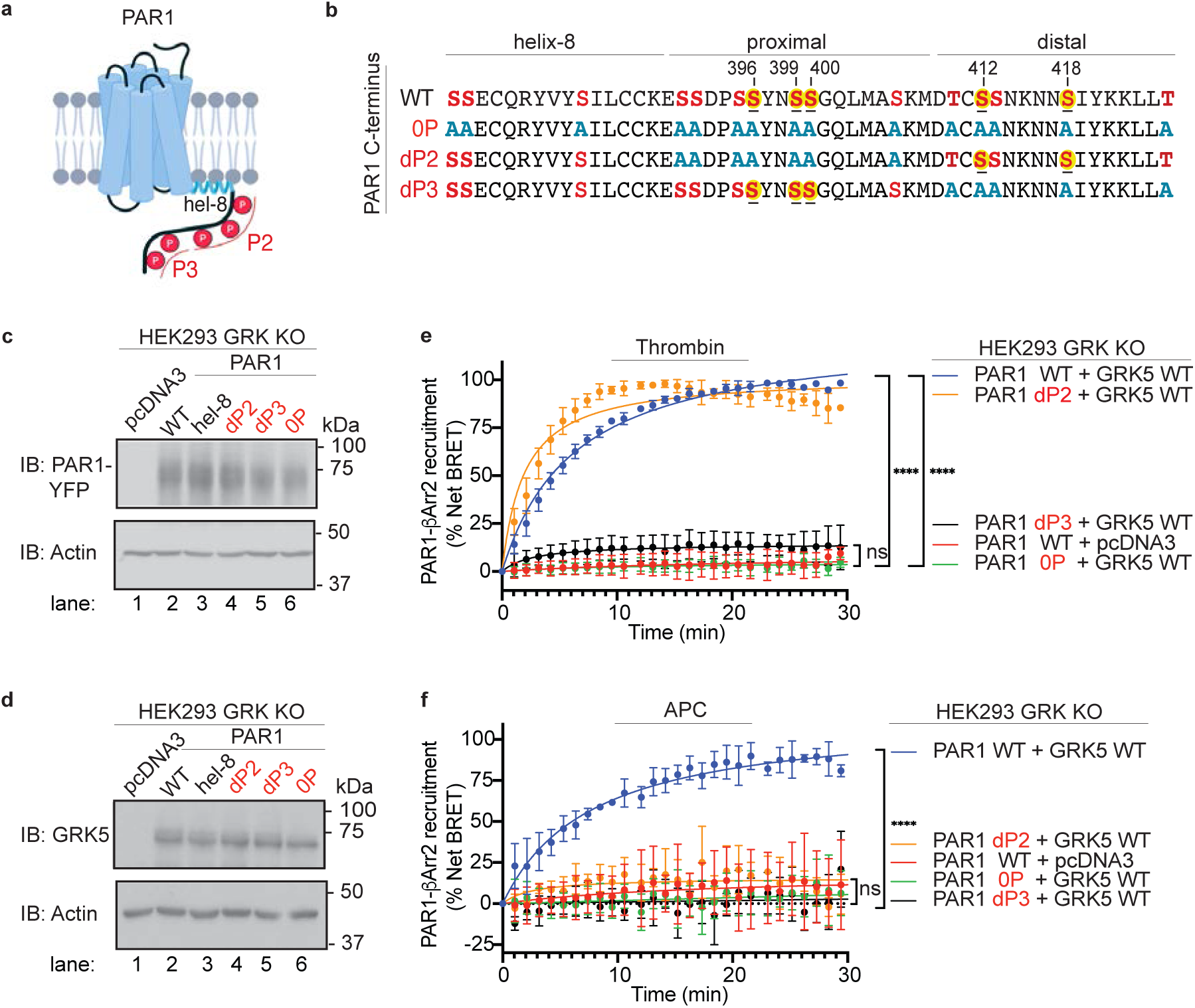
APC- *versus* thrombin-induced βarr2 recruitment require distinct PAR1 C-terminal phosphorylation sites. **a**, Cartoon of PAR1 C-terminus helix-8 and regions of phosphorylation. **b**, PAR1 C-terminus residues including known phosphorylation sites are numbered with yellow highlights. The PAR1 0P phospho-deficient mutant, proximal dP2 and distal dP3 phospho-site mutants with alanine (A) substitutions at Ser or Thr residues are indicated. **c** and **d**, HEK293 CRISPR-Cas9 GRK knockout (KO) cells were transfected with PAR1 WT, 0P, dP2 or dP3 phospho-site mutants together with GRK5 WT or pcDNA3 and immunoblotted as indicated. Actin was used as a loading control. **e** and **f**, HEK293 GRK KO cells transfected with PAR1-YFP WT or mutants, EPCR-Halo, Rluc-βarr2 and GRK5 WT or pcDNA3 were stimulated with 1 nM thrombin or 20 nM APC and βarr2 recruiment was determined by BRET. The data (mean ± S.D.) from three independent replicates was analyzed by one-way ANOVA followed by Tukey’s multiple comparisons test (****, *p* < 0.0001; ns = not significant).

To examine whether phosphorylation is required for GRK5-dependent βarr2 recruitment to activated PAR1, we used BRET assays. HEK293 GRK KO cells co-expressing PAR1 WT or the phospho-deficient PAR1 0P mutant fused to YFP (Fig. 5c) together with Rluc-βarr2, EPCR and either GRK5 WT or pcDNA3 (Fig. 5d) were stimulated with thrombin or APC and PAR1-βarr2 net BRET was determined. GRK5 WT expression rescued β-arr2 recruitment to thrombin- and APC-activated PAR1 WT but not to the PAR1 0P mutant (Fig. 5e, f), indicating that PAR1 phosphorylation mediated by GRK5 is required for both thrombin- and APC-induced βarr2 recruitment. Next, we examined whether specific PAR1 C-terminus phosphorylation sites are required for βarr2 recruitment to thrombin- or APC-activated PAR1 using the PAR1 proximal (dP2) and distal (dP3) phospho-site mutants (Fig. 5b, c) ^35^. Thrombin-induced βarr2 recruitment was abolished in cells expressing the PAR1 distal dP3 phospho-site mutant (Fig. 5e) and retained in cells expressing the proximal dP2 phospho-site mutant (Fig. 5e), indicating that the distal phospho-sites are critical for thrombin-induced βarr2 recruitment. Similar results were observed in HEK293 parental cells where thrombin-induced βarr2 recruitment was substantially reduced in cells expressing the PAR1 distal dP3 mutant but not in the proximal dP2 phospho-site mutant (Fig. S1a, c). In contrast to thrombin, both dP2 and dP3 PAR1 phospho-site mutants failed to recruit βarr2 in response to APC (Fig. 5f), indicating that both proximal and distal sites of phosphorylation are required for βarr2 recruitment to APC-stimulated PAR1. APC-stimulated βarr2 recruitment was also similarly inhibited in HEK293 parental cells expressing the PAR1 0P, dP2 and dP3 mutants (Fig. S1b, c), indicating that the results are not attributable to a GRK KO cell-line specific effect. These results demonstrate distinct phosphorylation requirements for βarr2 association with thrombin- *versus* APC-activated PAR1 where βarr2 is rapidly recruited to thrombin-activated PAR1 with only distal C-terminus residues phosphorylated, whereas βarr2 association with APC-activated PAR1 requires phosphorylation of the proximal and distal regions of the C-terminus. This agrees with APC-activated PAR1-induced βarr2 recruitment dependence on GRK5 plasma membrane localization (Fig. 4) that likely enables GRK5 access to the Ser/Thr sites in the proximal C-terminus of PAR1.

### APC but not thrombin promotes core engagement between βarr2 and activated PAR1

β-arrestins bind to activated GPCRs through at least two distinct modes: one where the finger loop region of β-arrestin inserts into the receptor transmembrane core ^36, 37, 38^ (the core-engaged configuration, Fig. 6a) and another exclusively mediated by the receptor’s phosphorylated C-terminus (tail-hanging configuration, Fig. 6a). To experimentally determine how thrombin- and APC-activated PAR1 engage with βarr2, we compared the recruitment of βarr2 WT to a mutant lacking the FLR (dFLR) using BRET assays. HEK293 βarr1,2 CRISPR-Cas9 knockout cells ^39^ co-expressing PAR1-YFP, EPCR and similar amounts of either Nanoluciferase (Nluc)-βarr2 WT or Nluc-βarr2-dFLR mutant (Fig. 6b, c), were stimulated with thrombin or APC and the change in PAR1-βarr2 net BRET was determined. Thrombin stimulated a significant increase in both βarr2 WT and dFLR mutant recruitment to activated PAR1 (Fig. 6d), suggesting that the FLR-mediated core engagement is dispensable for their interaction, and consistent with a tail-hanging configuration of βarr2. In contrast, APC-induced a robust and significant increase in βarr2 WT recruitment (Fig. 6e) but failed to promote recruitment of the βarr2 dFLR mutant (Fig. 6e). Since thrombin-activated PAR1 is desensitized by βarr1 ^14, 15^, we examined whether the FLR was required for βarr1 recruitment to PAR1 using HEK293 cells expressing Nluc-βarr1-WT and - dFLR mutant (Fig. 6f, g). In contrast to βarr2 dFLR, thrombin-induced recruitment of βarr1 dFLR to PAR1 was severely compromised (Fig. 6h), indicating that unlike βarr2, βarr1 engages with the transmembrane core of thrombin-activated PAR1.

**Fig. 6.**
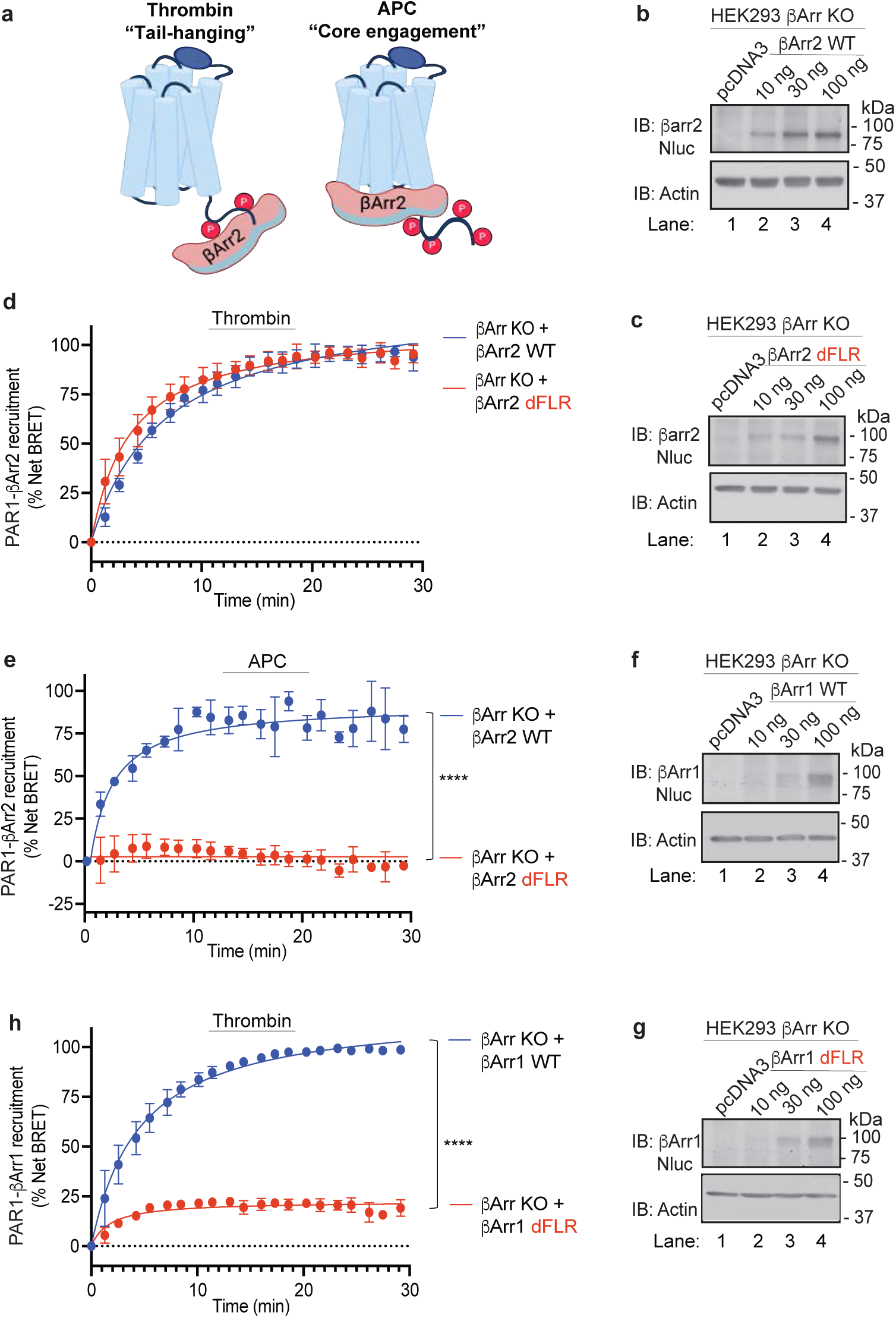
Distinct βarr2 determinants are required for APC- *versus* thrombin-induced βarr2 recruitment to PAR1. **a**, Cartoon of GPCR-βarr2 binding modes showing the tail-hanging and core engagment configurations. **b** and **c**, HEK293 CRISPR-Cas9 β-arrestin-1,2 knockout (KO) cells transfected with PAR1-YFP and EPCR-Halo and either pcDNA3 or varying amounts of Nluc-βarr2 WT or Nluc-βarr2 finger loop region (dFLR) mutant were immunoblotted as indicated. **d**and **e**, HEK293 CRISPR-Cas9 βarr1,2 KO cells transfected with PAR1-YFP, EPCR-Halo and a 100 ng of pcDNA3, Nluc-βarr2 WT or Nluc-βarr2 finger loop region (dFLR) mutant were stimulated with either 1 nM thrombin or 20 nM APC and βarr2 recruitment determined by BRET. The data (mean ± S.D.) from three independent repetitions were analyzed by one-way ANOVA followed by Tukey’s multiple comparisons test (****, *p* < 0.0001). **f** and **g**, HEK293 βarr1,2 KO cells transfected with PAR1-YFP, EPCR-Halo and either pcDNA3 or varying amounts of Nluc-βarr1 WT or dFLR mutant were immunoblotted as indicated. **h**, HEK293 βarr1,2 KO cells transfected with PAR1-YFP and EPCR-Halo and pcDNA3, Nluc-βarr1 WT or dFLR mutant were stimulated 1 nM thrombin and recruitment of βarr1 was determined by BRET. The data (mean ± S.D.) from three independent experiments was analyzed by one-way ANOVA followed by Tukey’s multiple comparisons test (****, *p* < 0.0001).

### Structural predictions of thrombin- and APC-activated PAR1 in complex with Gq-βarr2 or βarr2 only

Our results are consistent with thrombin-activated and not APC-activated PAR1 forming multimeric complexes simultaneously with the core-engaged heterotrimeric G protein and a tail-hanging βarr2. To reveal the structural basis for such complexes, we generated 100 models of thrombin-activated PAR1 with distinct phospho-sites (residues 42-425 with pT410, pS412, pS413 and pS418) and APC-activated PAR1 (residues 47-425 with pS391, pS392, pS395, pS396, pS399, pS400, pT410, pS412, pS413 and pS418) in complex with both Gq and βarr2 using AlphaFold 3. The models of thrombin-activated PAR1 Gq-βarr2 complex featured high conformational consistency across the ensemble, with the Gαq subunit C-terminal α-helix inserted in the receptor transmembrane core and the βarr2 N-domain bound to the distal C-terminus phospho-sites in a tail-hanging mode (Fig. 7a). The PAR1-Gq complexes featured low average predicted aligned error (PAE) and high interface predicted template modeling (ipTM) scores for the C-terminal helix of Gαq (Fig. 7c, d), indicative of high prediction confidence. However, AF3 failed to recognize the inability of APC-activated PAR1 to couple to Gq; therefore, it (incorrectly) predicted canonical core-engaged geometry for Gq and a tail-hanging geometry for βarr2 for both thrombin- and APC-activated PAR1 (Fig.7a, b). However, APC-activated PAR1-Gq-βarr2 ensemble was more divergent (Fig. 7b) and showed a higher PAE and lower ipTM score for the C-terminal helix of Gαq compared to thrombin-activated PAR1 (Fig. 7c, d), indicating lower prediction confidence. We also observed that both the average PAE and ipTM metrics for the C-terminal helix of Gαq in the multimeric complex with thrombin-activated PAR1 were more favorable than the corresponding metrics in the βarr2 core-engaged complex (Fig. 7c, d). By contrast, for APC-activated PAR1, the metrics of βarr2 core-engaged complex indicate higher confidence, compared to the multimeric Gq-containing complex (Fig. 7c, d). These findings are consistent with APC-activated PAR1 preference for βarr2 binding rather than G protein coupling ^11, 40^. This analysis supports our experimental data and demonstrates that thrombin- and APC-activated PAR1 conformations are distinct and enable preferential interaction with G proteins *versus* βarr2.

**Fig. 7.**
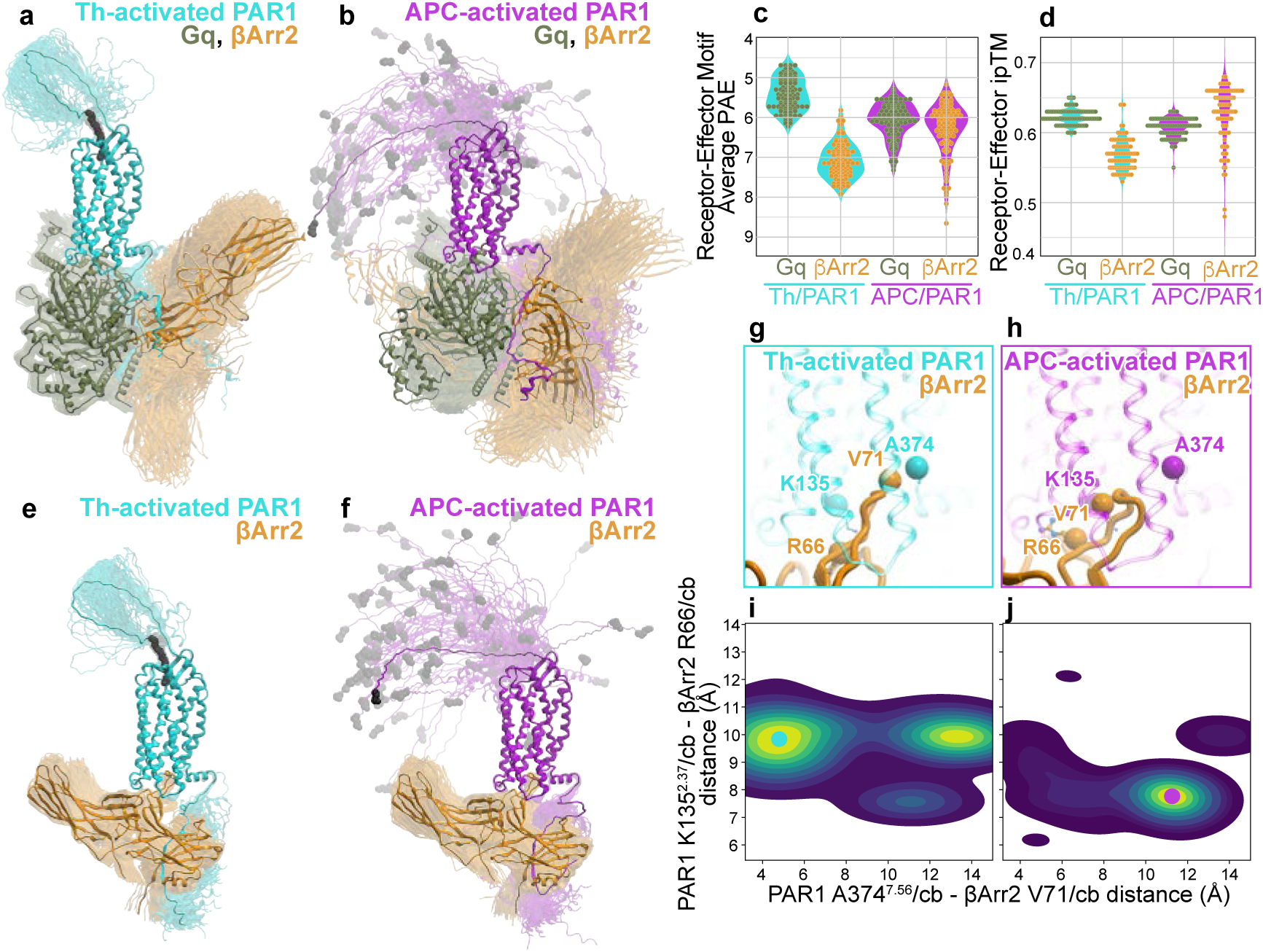
Structure prediction of thrombin- and APC-activated PAR1 bound to Gq-βarr2 or βarr2 only. Structural ensembles of 100 AlphaFold 3 models of **a**, thrombin-activated PAR1 bound to Gq-βarr2 (**a**) or βarr2 only (**e**) and APC-activated PAR1 bound to Gq-βarr2 (**b**) or βarr2 only (**f**). The receptors are shown as backbone ribbons, thrombin-activated PAR1 (*cyan*) with phosphorylated T410, S412, S413, and S418 residues (not shown) and APC-activated PAR1 (*magenta*) with phosphorylated S391, S392, S395, S396, S399, S400, pT410, S412, S413 and S418 residues. **c** and **d**, the distribution of average PAE and ipTM score values for the ensemble models of thrombin-activated and APC-activated PAR1 complexes with their effectors, Gq-βarr2 or βarr2. **g** and **h**, overlay of thrombin-activated PAR1 (*cyan*) or APC-activated PAR1 (*magenta*) with the finger loop region of βarr2 (*gold*) are shown as backbone ribbons and atoms of key residues used for calculating distances are shown as spheres and sticks. **i** and **j**, the distributions, across the model ensemble, of distances between key residues in PAR1 (K135, A374) and βarr2 (R66, V71) are shown as pseudo-colored two-dimensional density plots. The small, filled circles mark the representative conformations specific for thrombin-activated (*cyan*) and APC-activated PAR1 (*magenta*) with βarr2 finger loop region. Distances are measured in angstroms (Å).

To investigate the structural basis of favorable core-mediated coupling between βarr2 and APC-activated and not thrombin-activated PAR1, we constructed 100 models of only βarr2 complexes with activated PAR1, each containing distinct phospho-sites as described for Fig. 7a, b. When presented with these complex compositions, AF3 failed to predict the tail-handing mode for βarr2 with thrombin-activated PAR1 and instead predicted core-engaged model for all complexes (Fig. 7e, f). However, the geometry of the finger loop interaction with PAR1 was notably different in APC-activated complexes compared to thrombin-activated complexes (Fig. 7g, h). In APC-activated complexes, the preferred finger loop conformation was more compact (Fig. 7h), whereas in thrombin-activated complexes it was in an extended conformation (Fig. 7g). This is also demonstrated by the distances measured between key interaction residues in the PAR1 core region (K135^2.37^/cb and A374^7.56^/cb) and the FLR of βarr2 (R66/cb and V71/cb). In the thrombin-activated PAR1 ensemble, a substantial population of models have PAR1 A374 and βarr2 V71 in close proximity (Fig. 7i), whereas in APC-activated model ensemble, they are predominantly far apart (Fig. 7j). The predominant population in the APC-activated PAR1 ensemble also features shorter distances between PAR1 K135 and βarr2 R66, compared to thrombin-activated ensemble (Fig. 7i, j). Finally, compared with thrombin-activated PAR1, AF3 models of βarr2 complexes with APC-activated PAR1 have lower average PAE for βarr2 and higher ipTM scores, both indicative of higher confidence prediction (Fig.7c, d). Thus, despite being unable to predict the tail-handing mode for βarr2 complex with thrombin-activated PAR1, AF3 captured the conformational differences in the receptor transmembrane domain caused by differential N-terminal cleavage and C-terminal phosphorylation that was translated into distinct finger loop conformations and varying prediction confidence.

### Distinct βarr2 conformational changes induced by thrombin- *versus* APC-activated PAR1

β-arrestins adopt distinct activation states that are driven in part by the phosphorylation patterns on the C-termini of activated GPCRs. To examine whether thrombin and APC induce different βarr2 conformations, we used a series of intramolecular Nluc-βarr2 fluorescein arsenical hairpin (FlAsH)-based BRET biosensors ^41^. Each biosensor contains a tetracysteine motif that binds fluorescent arsenical (F) inserted at different positions in the N-domain or C-domain to monitor conformational changes induced by βarr2 recruitment to agonist-activated GPCRs (Fig. 8a, b) ^42^. To compare βarr2 conformational changes induced by thrombin and APC, HEK293 cells were transfected with individual Nluc-βarr2 N-domain F2, F3, F4 or F5 mutant biosensors or with individual Nluc-βarr2 C-domain F1, F7, F9 or F10 mutant biosensors and stimulated with saturating concentrations of thrombin or APC, after which net BRET was determined. Thrombin induced robust conformational changes in the βarr2 N-domain F2 and F5 biosensors and minimal conformational changes in F3 and F4 biosensors (Fig. 8c). By contrast, APC promoted modest conformational changes in all the βarr2 N-domain F2, F3, F4 and F5 biosensors (Fig. 8d). In studies with the βarr2 C-domain biosensors, thrombin caused a modest change in βarr2 C-domain F1, F7 and F10 mutant biosensors (Fig. 8f), but not in the F9 βarr2 biosensor. Similar modest conformational changes in βarr2 C-domain F1, F7, F9 and F10 mutant biosensor were observed with APC (Fig. 8f), except for F9 that retained sensitivity to APC stimulation, unlike thrombin (Fig. 8f, e). Together these results indicate that thrombin and APC induce distinct βarr2 conformational changes as summarized in the radar chart (Fig. 8g) and are predicted to translate into the distinct functional responses, particularly for APC-activated PAR1 βarr2 mediated cytoprotective signaling ^11, 12, 13^.

**Fig. 8.**
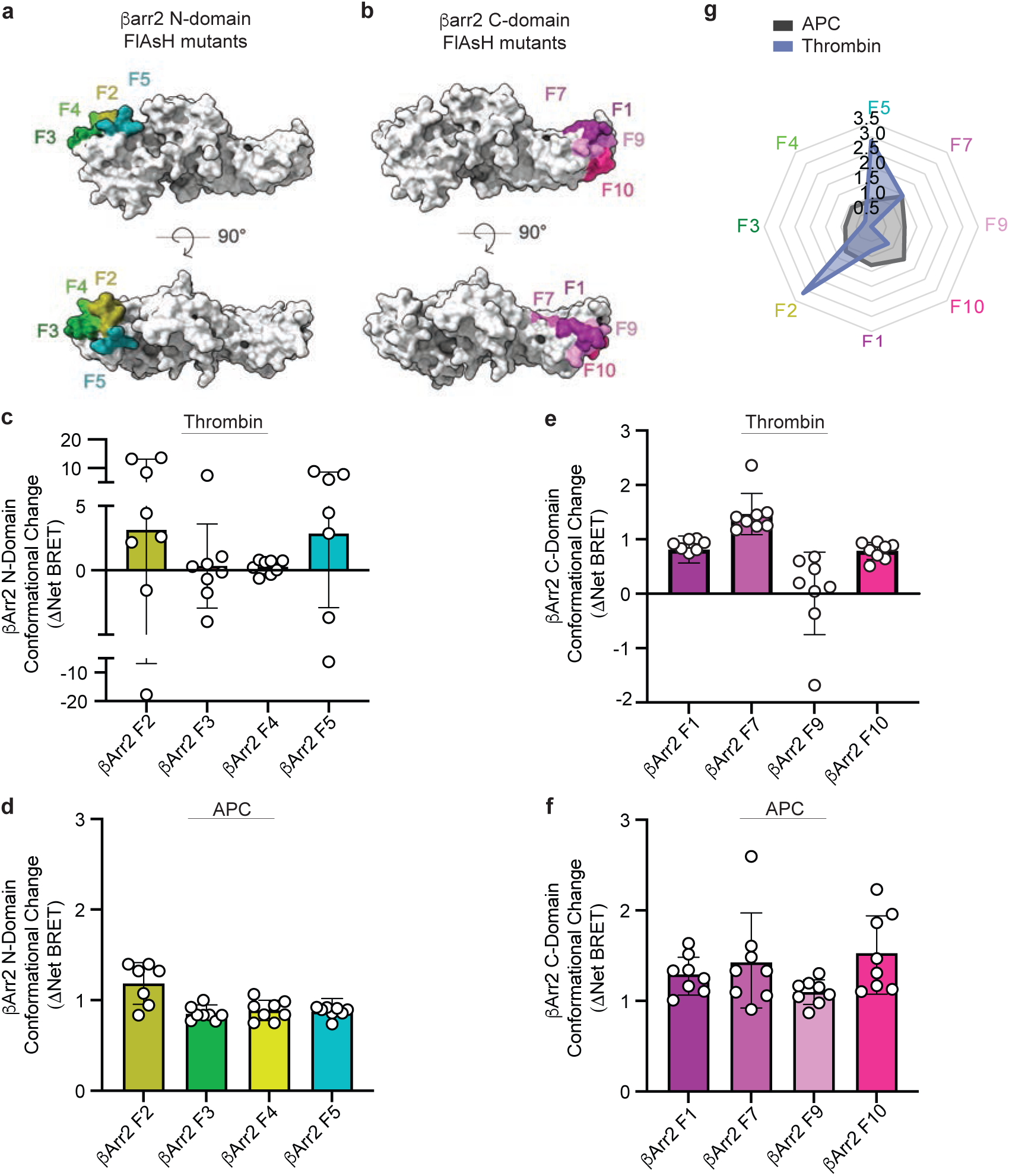
APC and thrombin induce distinct βarr2 conformational states. **a** and **b**, βarr2 inactive structure (PDB: 3P2D) surface projection of the N-domain F2, F3, F4 and F5 FlAsH-binding motifs (green palette colors) and C-domain F1, F7, F9 and F10 FlAsH-binding motifs (pink palette colors) of the NLuc-βarr2 FlAsH BRET. HEK293 cells transfected with untagged wildtype PAR1, EPCR-Halo and one of the individual NLuc-β-arr2 FlAsH N-domain or C-domain BRET biosensors were labeled with FlAsH reagent and stimulated with either 1 nM thrombin (**c** and **e**) or 20 nM APC (**d** and **f**) and net BRET determined. Data (mean ± S.D.) are from seven or eight independent replicates. **g**, Radar chart representation of APC-*versus* thrombin-induced changes in the βarr2 conformational biosensors shown in panels **c, d, e** and **f**.

## Discussion

In this study combined computational and experimental approaches were taken to determine the underlying mechanism of PAR1 biased signaling. Here, we report that PAR1 biased signaling is driven by different activated PAR1 conformational states but regulated by the same GRK5 isoform. Nonetheless, PAR1 biased agonism requires different GRK5 and βarr2 determinants to enable adoption of distinct βarr2 conformational states that is determined in part by differential requirement for GRK5 membrane anchoring. We also demonstrate that thrombin- *versus* APC-activated PAR1 recruitment of βarr2 utilizes different receptor C-terminus phosphorylation sites and different βarr2 structural elements. In support of our findings, AlphaFold 3 computational modeling further predicts that thrombin-activated PAR1 preferentially engages Gq-βarr2, whereas APC-activated PAR1 favors βarr2 engagement without the Gq protein in support of the above findings.

The mechanisms that specify which GRK isoform regulates a given GPCR are not clear. In human cultured endothelial cells, we found that GRK5 is the most highly expressed isoform and functions as the key regulator of both thrombin- and APC-activated PAR1 signaling. GRK5 was previously shown to regulate thrombin/PAR1 desensitization in endothelial cells ^43^. In contrast, in human platelets, PAR1 and the second thrombin receptor PAR4 are regulated by both GRK5 and GRK6 ^44^, although GRK6 is the most abundant isoform ^45^ in this cell type. In addition to high expression, the accessibility and conformation of activated PAR1 are likely important for specifying GRK5 isoform function. Unlike GRK2 and GRK3, GRK5 is primarily localized at the plasma membrane via a C-terminus amphipathic helix. In our study, expression of a plasma membrane deficient GRK5 4A amphipathic helix mutant in GRK KO cells failed to restore APC-induced βarr2 recruitment, whereas thrombin-induced βarr2 recruitment was partially rescued. Thus, membrane localization of GRK5 may increase prolonged access to APC-activated PAR1 to allow sufficient phosphorylation of proximal and distal sites necessary for βarr2 recruitment. In contrast, thrombin-activated PAR1-Gq signaling is likely to cause rapid release of GRK5 from the plasma membrane to the cytosol by inducing Ca^2+^-mediated calmodulin activation ^46^, thereby limiting access and phosphorylation of PAR1. Unlike thrombin/PAR1 rapid amplification of G protein signaling, APC-induced PAR1-βarr2 signaling onset is slow, prolonged and dependent on caveolae plasma membrane microdomains ^11, 40, 47^. GRK5 contains an N-terminal caveolin binding motif, binds caveolin-1 and localizes to caveolae ^48^. Thus, GRK5 membrane localization may also increase accessibility to PAR1 by concentrating these molecules in caveolae. However, the regulation of GPCR signaling by GRK5 localized to caveolae remains to be investigated.

The barcode hypothesis states that different GPCR phosphorylation patterns modulate β- arrestin function. Several studies have shown that GRK2/3 and GRK5/6 can phosphorylate the same GPCR at different sites to confer specific β-arrestin-driven functions such as internalization or signaling ^49, 50, 51^. Other studies reported that GPCR biased agonists utilize different GRKs to initiate distinct β-arrestin-mediated responses. The β2-adrenergic receptor (β2AR) biased agonist carvedilol recruits GRK5/6 more efficiently over other GRKs and promotes a different phosphorylation pattern and βarr2-elicited functions ^49, 52^. Utilization of GRK5/6 over GRK2/3 by the angiotensin II type-1 receptor (AT1R) biased agonist TRV027 also preferentially induces βarr2-dependent responses ^31^. In contrast to classic biased agonists, we found that GRK5 is utilized by both PAR1 biased agonists but utilizes different C-terminus phosphorylation patterns to enable βarr2 recruitment and distinct conformations.

β-arrestins are known to associate with agonist-activated GPCRs in at least two different configurations including the tail-hanging mode, where β-arrestin binds to the receptor C-terminus phospho-sites and the core-engaged mode, where β-arrestin FLR inserts into the receptor transmembrane core ^36, 37, 38^. We found that thrombin- and APC-activated PAR1 engage with βarr2 using different binding modes. APC induced βarr2 recruitment requires the FLR structural element indicating core engagement, whereas the FLR is dispensable for thrombin-stimulated βarr2 recruitment suggesting a tail-hanging mode. The different PAR1-βarr2 binding configurations are likely enabled by the distinct C-terminus phosphorylation patterns required. APC/PAR1-βarr2 interaction requires phospho-sites spanning the proximal P2 and distal P3 regions, whereas only the distal P3 phospho-sites are necessary for thrombin/PAR1-βarr2 engagement. AlphaFold 3 modeling indicates high confidence for the APC-activated PAR1-βarr2 complex with core engagement, whereas the thrombin-activated PAR1-Gq-βarr2 is more structurally aligned with the tail-hanging conformation. In contrast, βarr1 recruitment induced by thrombin-activated PAR1 is dependent on the FLR structural element and was shown to require the P3 distal phosphorylation sites of PAR1 ^53^, similar to βarr2. Thus, β-arrestin isoforms are not redundant and can engage the same agonist-activated GPCR using the same phosphorylation sites, yet adopt distinct conformations, a phenomenon that may rely on isoform specific intrinsic features.

The different GPCR-β-arrestin binding modes confer distinct functions. The current paradigm postulates that GPCR-β-arrestin tail-hanging mode facilitates internalization and continued G protein signaling, whereas core engagement promotes desensitization and termination of G protein signaling ^1, 54^. Our study shows that thrombin- and APC-activated PAR1 engage with distinct βarr1 and βarr2 configurations that have both similar and different functional implications compared to the established paradigm. In fibroblasts derived from β-arrestin knockout mice, we showed that βarr1 is critical for thrombin/PAR1 desensitization and neither βarr1 nor βarr2 are required for receptor internalization ^14, 15^. Rather, PAR1 internalization occurs through a clathrin adaptor protein-2 dependent pathway ^55, 56, 57^. Thus, thrombin-activated PAR1-βarr1 core engagement prevents G protein activation and is dispensable for receptor internalization, consistent with the prevailing paradigm. The function of thrombin/PAR1-βarr2 tail-hanging engagement is not known but this configuration is permissive of receptor association with other effectors ^36^ including simultaneous G protein coupling ^58, 59^, as is predicted by AlphaFold 3 modeling. Although βarr2 is required for APC/PAR1 biased signaling in various cell types and mouse models ^11, 13, 29, 60^, βarr2 fails to promote APC-activated PAR1 internalization. These findings are consistent with an APC-induced PAR1-βarr2 core engagement configuration that occludes G protein activation and elicits βarr2 signaling but is incapable of promoting receptor internalization. Currently, it is not known why the APC-activated βarr2 conformation fails to promote receptor internalization: a phenomenon that may be due to the receptor phosphorylation state, membrane microdomain localization and membrane lipid composition or other features inherent to βarr2.

In conclusion, we report a new regulatory mechanism for GPCR biased signaling that is mediated by GRK5. The GRK5 isoform promotes distinct βarr2 activated conformational states induced by biased agonists that is driven by different GPCR C-terminus phosphorylation patterns. We also discovered that the GRK5 amphipathetic helix and βarr2 FLR region, distinct structural elements, mediate the specificity of APC v*ersus* thrombin biased signaling. The novel regulatory mechanism for GPCR biased signaling is strongly supported by computational analysis and experimental validation.

## Methods

### AlphaFold 3 modeling

AlphaFold 3 ^25^ server was used to generate three-dimensional models of human PAR1 with varying phosphorylation patterns, with and without effector proteins.

The following amino-acid sequences and phosphorylation patterns were used for PAR1 (all based on UniProt entry P25116, PAR1_HUMAN):

PAR1(22-390) (mature unactivated PAR1 lacking the disordered C-terminus)

ARTRARRPES KATNATLDPR SFLLRNPNDK YEPFWEDEEK NESGLTEYRL VSINKSSPLQ KQLPAFISED ASGYLTSSWL TLFVPSVYTG VFVVSLPLNI MAIVVFILKM KVKKPAVVYM LHLATADVLF VSVLPFKISY YFSGSDWQFG SELCRFVTAA FYCNMYASIL LMTVISIDRF LAVVYPMQSL SWRTLGRASF TCLAIWALAI AGVVPLLLKE QTIQVPGLNI TTCHDVLNET LLEGYYAYYF SAFSAVFFFV PLIISTVCYV SIIRCLSSSA VANRSKKSRA LFLSAAVFCI FIICFGPTNV LLIAHYSFLS HTSTTEAAYF AYLLCVCVSS ISCCIDPLIY YYASSECQRY VYSILCCKE

PAR1(42-390) (thrombin-activated PAR1 lacking the disordered C-terminus)

SFLLRNPNDK YEPFWEDEEK NESGLTEYRL VSINKSSPLQ KQLPAFISED ASGYLTSSWL TLFVPSVYTG VFVVSLPLNI MAIVVFILKM KVKKPAVVYM LHLATADVLF VSVLPFKISY YFSGSDWQFG SELCRFVTAA FYCNMYASIL LMTVISIDRF LAVVYPMQSL SWRTLGRASF TCLAIWALAI AGVVPLLLKE QTIQVPGLNI TTCHDVLNET LLEGYYAYYF SAFSAVFFFV PLIISTVCYV SIIRCLSSSA VANRSKKSRA LFLSAAVFCI FIICFGPTNV LLIAHYSFLS HTSTTEAAYF AYLLCVCVSS ISCCIDPLIY YYASSECQRY VYSILCCKE

PAR1(47-390) (APC-cleaved PAR1 lacking the disordered C-terminus)

NPNDKYEPFW EDEEKNESGL TEYRLVSINK SSPLQKQLPA FISEDASGYL TSSWLTLFVP SVYTGVFVVS LPLNIMAIVV FILKMKVKKP AVVYMLHLAT ADVLFVSVLP FKISYYFSGS DWQFGSELCR FVTAAFYCNM YASILLMTVI SIDRFLAVVY PMQSLSWRTL GRASFTCLAI WALAIAGVVP LLLKEQTIQV PGLNITTCHD VLNETLLEGY YAYYFSAFSA VFFFVPLIIS TVCYVSIIRC LSSSAVANRS KKSRALFLSA AVFCIFIICF GPTNVLLIAH YSFLSHTSTT EAAYFAYLLC VCVSSISCCI DPLIYYYASS ECQRYVYSIL CCKE

PAR1(42-425) (full-length thrombin-activated PAR1)

SFLLRNPNDK YEPFWEDEEK NESGLTEYRL VSINKSSPLQ KQLPAFISED ASGYLTSSWL TLFVPSVYTG VFVVSLPLNI MAIVVFILKM KVKKPAVVYM LHLATADVLF VSVLPFKISY YFSGSDWQFG SELCRFVTAA FYCNMYASIL LMTVISIDRF LAVVYPMQSL SWRTLGRASF TCLAIWALAI AGVVPLLLKE QTIQVPGLNI TTCHDVLNET LLEGYYAYYF SAFSAVFFFV PLIISTVCYV SIIRCLSSSA VANRSKKSRA LFLSAAVFCI FIICFGPTNV LLIAHYSFLS HTSTTEAAYF AYLLCVCVSS ISCCIDPLIY YYASSECQRY VYSILCCKES SDPSSYNSSG QLMASKMDTC SSNLNNSIYK KLLT

PAR1(47-425) (full-length APC-cleaved PAR1)

NPNDKYEPFW EDEEKNESGL TEYRLVSINK SSPLQKQLPA FISEDASGYL TSSWLTLFVP SVYTGVFVVS LPLNIMAIVV FILKMKVKKP AVVYMLHLAT ADVLFVSVLP FKISYYFSGS DWQFGSELCR FVTAAFYCNM YASILLMTVI SIDRFLAVVY PMQSLSWRTL GRASFTCLAI WALAIAGVVP LLLKEQTIQV PGLNITTCHD VLNETLLEGY YAYYFSAFSA VFFFVPLIIS TVCYVSIIRC LSSSAVANRS KKSRALFLSA AVFCIFIICF GPTNVLLIAH YSFLSHTSTT EAAYFAYLLC VCVSSISCCI DPLIYYYASS ECQRYVYSIL CCKESSDPSS YNSSGQLMAS KMDTCSSNLN NSIYKKLLT

The following amino-acid sequences were used for effector proteins: Human βarr2(1-351) (based on UniProt P32121, ARRB2_HUMAN)

MGEKPGTRVF KKSSPNCKLT VYLGKRDFVD HLDKVDPVDG VVLVDPDYLK DRKVFVTLTC AFRYGREDLD VLGLSFRKDL FIATYQAFPP VPNPPRPPTR LQDRLLRKLG QHAHPFFFTI PQNLPCSVTL QPGPEDTGKA CGVDFEIRAF CAKSLEEKSH KRNSVRLVIR KVQFAPEKPG PQPSAETTRH FLMSDRSLHL EASLDKELYY HGEPLNVNVH VTNNSTKTVK KIKVSVRQYA DICLFSTAQY KCPVAQLEQD DQVSPSSTFC KVYTITPLLS DNREKRGLAL DGKLKHEDTN LASSTIVKEG ANKEVLGILV SYRVKVKLVV SRGGDVSVEL PFVLMHPKPH D

Human Gαq (1-354) (based on UniProt P50148, GNAQ_HUMAN)

MTLESIMACCLSEEAKEARRINDEIERQLRRDKRDARRELKLLLLGTGESGKSTFIKQMRIIHGSGYSDEDKRGFTKLVYQNIFTAMQAMIRAMDTLKIPYKYEHNKAHAQLVREVDVEKVSAFENPYVDAIKSLWNDPGIQECYDRRREYQLSDSTKYYLNDLDRVADPAYLPTQQDVLRVRVPTTGIIEYPFDLQSVIFRMVDVGGQRSERRKWIHCFENVTSIMFLVALSEYDQVLVESDNENRMEESKALFRTIITYPWFQNSSVILFLNKKDLLEEKIMYSHLVDYFPEYDGPQRDAQAAREFILKMFVDLNPDSDKIIYSHFTCATDTENIRFVFAAVKDTILQLNLK

Human Gβ1 (1-340) (UniProt P62873, GBB1_HUMAN)

MSELDQLRQEAEQLKNQIRDARKACADATLSQITNNIDPVGRIQMRTRRTLRGHLAKIYAMHWGTDSRLLVSASQDGKLIIWDSYTTNKVHAIPLRSSWVMTCAYAPSGNYVACGGLDNICSIYNLKTREGNVRVSRELAGHTGYLSCCRFLDDNQIVTSSGDTTCALWDIETGQQTTTFTGHTGDVMSLSLAPDTRLFVSGACDASAKLWDVREGMCRQTFTGHESDINAICFFPNGNAFATGSDDATCRLFDLRADQELMTYSHDNIICGITSVSFSKSGRLLLAGYDDFNCNVWDALKADRAGVLAGHDNRVSCLGVTDDGMAVATGSWDSFLKIWN

Human Gγ (1-69) (UniProt P59768, GBG2_HUMAN)

MASNNTASIAQARKLVEQLKMEANIDRIKVSKAAADLMAYCEAHAKEDPLLTPVPASENPFREKK FFCA

One hundred models (20 random seeds with 5 models per seed) were built for each of the following proteins and complexes:

- Uncomplexed PAR1(22-390)
- Uncomplexed PAR1(42-390)
- Uncomplexed PAR1(47-390)
- PAR1(amino acids 42-425, containing pT410, pS412, pS413, pS418) with βarr2
- PAR1(amino acids 47-425, containing pS391, pS392, pS395, pS396, pS399, pS400, pT410, pS412, pS413, pS418) with βarr2
- PAR1(amino acids 42-425, containing pT410, pS412, pS413, pS418) with Gαq, Gβ1, Gγ2, and βarr2
- PAR1(amino acids 47-425, containing pS391, pS392, pS395, pS396, pS399, pS400, pT410, pS412, pS413, pS418) with Gαq, Gβ1, Gγ2, and βarr2

For uncomplexed receptor models, the removal of the receptor C-terminus (amino acids 391-425) prevented the conformational bias due to its frequently predicted insertion into the effector binding surface and thus enabled the studies of conformational preferences of PAR1 TM domain as a function of N-terminal proteolytic cleavage. For models with βarr2, the removal of the βarr2 C-terminus (amino acids 352-409) prevented the self-inhibited βarr2 conformation where its C-terminus blocks the N-lobe interface and enabled studies of binding preferences for the differentially phosphorylated PAR1 C-terminus.

The resulting complexes were superimposed by the receptor TM domain, visualized, and analyzed in ICM v3.9-3a (Molsoft LLC, San Diego, CA). Inter-and intramolecular distances were measured using python library *MDAnalysis* ^61^, 2D pseudo-color histograms were plotted with *matplotlib* ^62^. Confidence metrics (pLDDT, PAE, ipTM) were extracted from the json files generated by the AlphaFold 3 server. Average effector motif PAE values were calculated for amino acids 64-77 of βarr2 and amino acids 332-359 of Gαq. Violin plots were built in R using *ggplot2*^63^.

### Cell Culture

HUVEC-derived endothelial EA.hy926 cells (ATCC #CRL-2922) were maintained in 10% fetal bovine serum (FBS) (Gibco 10437-028) in Dulbecco’s Modified Eagle Medium (DMEM) (Corning 10-013-CV) supplemented with fresh 20% preconditioned media every 2-3 days, grown at 37°C in 8% CO_2_ and used up to passage 8 ^12^. Primary HUVECs (Lonza, #C2519A) were maintained in endothelial cell growth medium-2 (Lonza, #CC-3162) and media was changed every 2 days, grown at 37°C in 5% CO_2_ and used up to passage 6. HEK293A parental cells, HEK293A CRISPR-Cas9 β-arrestin-1,2 KO cells, and HEK293A GRK 2,3,5,6 KO cells were obtained from Dr. Asuka Inoue (Tohoku University) ^64^. Cells were maintained in 10% FBS in DMEM supplemented with fresh media every 2 to 3 days, grown at 37°C in 5% CO_2_ and used up to passage 10. HeLa-PAR1 cells were generated as previously described ^55^ and grown in DMEM supplemented with 10% FBS and 250 μg/ml hygromycin B and used up to passage 10.

### Antibodies and Reagents

Anti-GRK4-6 A16/17 (#05-466), anti-GRK2/3 C5/1.1 (#05-465), anti-Renilla Luciferase (#MAB4400), and anti-β-Actin AC-74 (#A5316) were from Millipore Sigma. Polyclonal anti-GRK5 (#PA5-96262) was from Invitrogen. Anti-phospho-p38 MAPK T180/Y182 (#4411), anti-p38 MAPK polyclonal (#9212), anti-phospho-Akt-S473 D9E (#4060), and anti-Akt polyclonal (#9272) were from Cell Signaling Technology. Anti-GFP c3H9 (#3h9) was from ChromoTek. Anti-mouse (#170–6516) and anti-rabbit (#170–6515) horseradish peroxidase–conjugated antibodies were from Bio-Rad and anti-mouse Alexa-488 Fluor (#A-11001) from Invitrogen. DAPI (#D-1306) was purchased from ThermoFisher Scientific. α-thrombin (#HT 1002a) was from Enzyme Research Laboratory, APC (#HCAPC-0080) from Prolytix, Vorapaxar (#1755) from Axon Medchem, and Coelentrazine H (#1011-1) from Biotium.

### Plasmids

GRK5 WT, K215R, and 4A mutant pcDNA3 plasmids were from Dr. Philip B. Wedegaertner (Thomas Jefferson University). PAR1-YFP, EPCR-Halo, RLuc-βarr2, PAR1 C-terminus phospho-site mutants 0P, P2, and P3 fused to YFP were generated by Gibson assembly homologous recombination (Gibson Assembly® Master Mix, New England Biolabs) followed by whole plasmid sequencing. The Nluc-βarr1 and -βarr2 WT and dFLR mutant plasmids were from Dr. Asuka Inoue (Tohoku University) and Nluc-βarr2 FlAsH variants plasmids were from Dr. Carsten Hoffmann (University Hospital Jena).

### Quantitative reverse-transcriptase polymerase chain reaction (qRT-PCR)

Endothelial cells were seeded in a 6-well plate at 3.2 x 10^5^ cells per well and grown to confluency. RNA was extracted using the Direct-zol RNA Miniprep Plus Kit (#R2072, Zymo Research) and used to generate complementary DNA (cDNA) according to the manufacturer’s instructions. RNA was quantified and cDNA synthesized from 1 μg RNA using SuperScript IV VILO Master Mix with ezDNase enzyme kit (#111766050, Thermo Fisher Scientific). qRT-PCR was performed with TaqMan Fast Advanced Master Mix (#4444964, Thermo Fisher Scientific) and TaqMan Gene Expression Probes GRK2 (#Hs00176395), GRK3 (#Hs00178266), GRK5 (#Hs00992173), GRK6 (#Hs00357776), and 18S (#Hs03003631_g1) using a QuantStudio 3 Real-Time PCR System (Thermo Fisher Scientific). GRK mRNA transcript levels were normalized to 18S expression. The differences in expression relative to GRK5 were then determined using the 2-^ΔΔ^Ct method. Control reactions without cDNA for each probe were conducted in every assay to ensure specificity of the reactions.

### Small interfering (si) RNA transfections

Endothelial cells were seeded in a 12-well plate at 2.5 × 10^5^ cells per well, grown overnight, and transfected with 25 nM GRK2 #1 siRNA (5′-CCGGGAGATCTTCGACTCATA-3′ Qiagen #SI00287378), 25 nM GRK5 #5 siRNA (5′-AGCGTCATAACT AGAACTGAA-3′ (Qiagen, #SI00287770) or 25 nM non-specific siRNA (AllStars Negative Control non-specific siRNA Qiagen, #1027281) using the TransIT-X2 System (Mirus, #MIR 600) according to the manufacturer’s instructions. Whole cell lysates were collected 48 h post transfection.

### Signaling assays and immunoblotting

After siRNA transfection, endothelial cells were serum-starved overnight in 0.4% FBS-DMEM. Cells were then incubated in serum-free DMEM containing 10 mM HEPES, 1 mM CaCl_2_, and 1 mg/mL bovine serum albumin (BSA) for 1 h prior to incubation with thrombin (10 nM) or APC (20 nM) at 37°C. After agonist stimulation, cells were lysed in 2x Laemmli sample buffer (LSB) containing 200 mM dithiothreitol (DTT), heated for 5 min at 95°C, resolved by SDS-PAGE and immunoblotted.

HEK293 cells were harvested with Triton lysis buffer (50 mM Tris pH 7.4, 100 mM NaCl, 5 mM EDTA, 1% v/v TritonX-100, 50 mM NaF, and 10 mm NaPP) containing protease inhibitors including 1 µg/ml of leupeptin, aprotinin, trypsin protease inhibitor or pepstatin, benzamidine 100 µg/ml, PMSF 100 µg/ml and quantified by bicinchoninic acid (BCA) protein assay (Thermo Fisher Scientific). Equivalent amounts of cell lysates were diluted in 2x LSB with 200 mM DTT, heated for 5 min at 95°C, resolved by SDS-PAGE, transferred to the PVDF membrane and immunoblotted. Immunoblots were quantified by densitometry using NIH Image J software.

### Immunofluorescence confocal microscopy

HeLa cells were seeded at 1.0 x 10^5^ per cover slip transfected with GRK5 WT, K215R, 4A mutant or pcDNA3. Cells were fixed with 4% paraformaldehyde, permeabilized with 0.1% Triton-X100 and labeled with monoclonal anti-GRK5 antibody (at 1: 500) for 1 h on ice. Cells were then incubated with anti-mouse Alexa-488 diluted to 1:750 and DAPI at 1 μg/ml in 0.03% BSA, 0.01% Triton-X 100, and 0.01% normal goat serum at room temperature for 1 h. Slides were mounted using ProLong Gold Antifade Mountant (Invitrogen, #P10144). Confocal images were acquired with an Olympus IX81 spinning-disk microscope equipped with a CoolSNAP HQ2 CCD camera (Andor) and 63× Plan Apo objective (1.4 NA) with appropriate excitation-emission filters using Metamorph software. Line scan analysis was performed using NIH Image J software.

### Bioluminescence resonance energy transfer (BRET) assays

HEK293 cells were seeded in a 6-well plate at 4.5 x 10^5^, grown overnight, and transfected with PAR1 WT, 0P, dP2 or dP3 mutants fused to YFP, APC co-receptor EPCR-Halo and, RLuc-βarr2 wildtype or pcDNA3 diluted in Opti-Mem and transfected with polyethylenimine (PEI) at a 1:3 ratio. In other BRET experiments, GRK5 WT, 4A or K215R and Nluc-βarr2- or Nluc-βarr1WT and dLR mutants were combined with wildtype PAR1-YFP and EPCR-Halo diluted in Opti-Mem and transfected with PEI. After 24 h of transfection, cells were collected and re-seeded into poly-D-lysine coated 96-well plate at 3 x 10^4^ cells per well and grown overnight. Cells were washed with PBS and serum-starved for 1 h using a 1:1 equal mixture of DMEM without phenol red (Gibco, #31053–028) combined with PBS. In some experiments, cells were pretreated with 10 μM vorapaxar for 30 min during starvation. After starvation, cells were preincubated with 5 µM of Coelenterazine H for 5 min followed by the addition of 1 nM thrombin or 20 nM APC and BRET measurements were taken at 37°C over time. All BRET measurements were performed with a Berthold TriStar LB941 multimode plate reader using MicroWIN 2010 software (Berthold Biotechnologies) using two filter settings: 480 nm for Rluc and 530 nm for YFP. The BRET signal was calculated as the emission at 530 nm divided by the emission at 480 nm. The BRET signals were normalized to basal BRET ratios and expressed as the percent over basal.

FlAsH BRET assays were performed in HEK293 cells, seeded, grown and transfected with wildtype PAR1-YFP, EPCR-Halo and the Nluc-βarr2 N-domain or C-domain FlAsH constructs ^41^ as described in BRET assays above. Cells were serum starved and incubated with 2 μM FlAsH-EDT2 (#T34561, ThermoFisher) for 30 min, washed with BAL (25 mM dimercaprol in 10 mM HEPES buffer, pH 7.3) for 5 min, and serum starved for an additional 30 min. After starvation, cells were preincubated with 5 µM of Coelenterazine H for 3 min followed by the addition of 1 nM thrombin or 20 nM APC and BRET measurements were taken for 5 min at 37°C and net BRET determined as described above.

### Statistical analysis

Data were analyzed using Prism 10.2 statistical software (GraphPad Software, La Jolla, CA) and Microsoft Excel. Statistical test analysis methods are indicated in the figure legends.

### Software

BioRender was used to create cartoons. Figures were created in Adobe Illustrator. βarr2 FlAsH mutants cartoons were created using the University of California, San Francisco Chimera X (version 1.9).

### Data availability

All study data are included in the article and available upon request.

## Acknowledgments and funding sources

We thank all members of the Trejo lab for advice and guidance and Hannah Higa for assistance with Figure illustrations. This work was supported by NIH/ NHLBI R01HL163931 (J.T.), NIH/ NHLBI T32HL007444 (M.R.G., C.A.B.), NIH/ NIGMS K12GM068524 (M.R.G), American Heart Association Postdoctoral Fellowship #25POST1369723 (L.O.C.) and NIH/ NIAID R21 AI156662 and R01AI161880 (I. K.)

## Author Contributions

MRG, LOC, YL, HQ, CAB, CB, IK and JT designed research; MRG, LOC, YL, HQ, CB and CAB performed research; MRG, LOC, HQ, CB, IK and JT analyzed data; MRG, LOC, CB, IK and JT wrote and revised the paper.

